# Molecular Characterisation Of Urine And Vaginal Samples And Their Antibiotic Susceptibility Pattern Associated With β-Lactamases Resistance Gene

**DOI:** 10.1101/2022.03.18.484962

**Authors:** Sonia Sethi, Rinki mishra, Vaishnavi gupta, Sandeep Shrivastava

**Affiliations:** Dr. B. Lal Institute of Biotechnology, Malviya Industrial Area, Malviya Nagar, Jaipur-302017

**Keywords:** Antibiotic sensitivity, Enterobacteriaceae, ESBL, CTX-M, TEM gene

## Abstract

Infection or inflammation of the vagina is called vaginitis. The two most common causes of vaginitis are bacterial vaginitis and Candida vaginitis. The causative organisms may be sexually transmitted or endogenous, usually by *Gardnerella vaginalis* and *Mycoplasma hominis* in combination with other anaerobes. Urinary tract infections (UTI) refer to the presence of microbial pathogens within the urinary tract and are caused by a range of pathogens but most commonly by *Escherichia coli, Klebsiella pneumoniae* etc. Highly increase in antibiotic-resistant among uropathogens, including ESBL producing E.coli, AmpC β-lactamase or carbapenemase-producing Enterobacteriaceae (e.g., New Delhi metallo-β-lactamase [NDM]) are being reported among health care unit. Carbapenem-resistant Enterobacteriaceae (CRE) is a growing concern worldwide and a major challenge by the WHO 2014.

The aim of this study was to determine the occurrence of CTX-M gene and TEM gene and the possible role of ESBL in resistance to antibiotic. A study was conducted on a total of 100 urine and vaginal samples from Dr. B. Lal clinical laboratory and was cultured on Nutrient Agar and MacConkey agar for the detection of etiologic agents. Bacteria isolation, identification & biochemical characterization was done. Kirby-Bauer Disk Diffusion (KBDS) method was used for detection of ESBL-producing strains. A disc of Cefotaxime (30 µg) and Clavulanic acid (30 µg) were placed at a distance of 20 mm on Muller Hilton agar plate along with Imipenem(30µg), Piperacillin/Tazobactam(10µg) and Ceftazidime(30µg). The zone of inhibition generally greater than 20mm was considered as ESBL producer. Total 14 isolates were identified as ESBL producer. In order to find out the plasmid profile of these isolates, plasmid DNA isolation was done. Isolates was found to have one or more than one plasmid gene responsible for the ESBL resistance. This finding concludes that CTX-M gene is most common gene in ESBL producing isolates. It is highly recommended that antibiotic prescription should be monitored according to guideline. The prevalence of ESBL is increasing day by day and necessary steps should be taken to prevent the spread and emergence of resistance.

## Introduction

In recent times, because of augmentation of bacterial resistance to various antibiotics, the management of irresistible diseases due to harmful pathogenic bacteria has become a challenge [1]. In medical system resistance for antibiotics has become a foremost issue and mainly in urinary tract infections. Although, UTI is curable, but nowadays becoming burdensome to manage the disease due to antibiotic resistance, especially in the *Enterobacteriaceae* family [2]. The traditional mechanism of resistance among the *Enterobacteriaceae* family is the exhibition of hydrolytic enzymes, the “*β*-lactamases” [3]. Increase in prevalence of *β*-lactamases causes complications in UTIs producing uropathogens [4]. ESBL producing organisms infections are linked with high medical expenditure, mortality and morbidity. ESBLs occur principally among *Escherichia coli* and *Klebsiella* species which are plasmid mediated and genes transfer horizontally between GNB members [5]. Out of all gram-negative bacteria that produce *β*-lactamases are of major concern due to their ability to spread globally and the assessable inadequate curable alternative because of multiple resistance genes and the enzymes’ which are associated with resistance to various non-beta-lactam antibiotics [6]. ESBLs are Class A β lactamases that are forbidden in vitro by β-lactamase inhibitors such as clavulanic acid and cannot hydrolyse cefoxitin or cefotetan i.e. cephamycins or imipenem and meropenem i.e. carbapenems but can hydrolyze penicillin, oxyimino-cephalosporins, and monobactams [7]. Studies of the antimicrobial susceptibility of GNB revealed that there is resistance augmentation due to hyperproduction of ESBL and AmpC enzymes over time. This phenomenon was observed among *E. coli, K. pneumoniae* and other GNB [8].

ESBLs have different genotypes and the most common are the SHV, TEM, and CTX-M types. Other types include VEB, PER, BEL-1, BES-1, SFO-1, TLA, and IBC [9]. Detection of Beta Lactamases resistant strains proves to be a guide in treatment using antibiotics and also minimizes spreading of infections [10].

## 2. Materials and Methods

### 2.1 Study design and population

This study was conducted at Dr. B. Lal Institute of Biotechnology, Jaipur (Rajasthan). A total of 100 samples of bacterial isolates were isolated and identified from different samples i.e. Urine and vaginal processed at Dr. B. Lal Clinical Laboratory Pvt. Ltd.

### 2.2 Processing of Samples

All Mid Stream Urine samples were cultured on routine culture media by semi-quantitative method as described in World Health Organization (WHO) manual [11]. In short, 1μL of urine was inoculated on MacConkey agar plate (Hi Media Laboratories Pvt. Ltd., India) by streaking using calibrated loop, and incubated aerobically for 18-48hrs at 37ᵒC. Samples that did not yield significant bacterial growth, those that had multiple organisms and samples with suspected contamination as per lab report, were excluded from the study. Growth of 100 colonies or more, *i.e*. 10^5^ colony forming units (CFU)/mL urine, was considered as culture positive. Isolation and identification of isolates were done following their morphology in Gram’s staining, cultural characteristics and biochemical properties, mainly IMViC (Indole test, Methyl red test, Voges-proskauer test and Citrate test) as per the Manual of Clinical Microbiology [12].

### 2.3 Antimicrobial Susceptibility Testing

Antibiotic susceptibility testing of all isolates was performed by Kirby-Bauer’s disc diffusion method and interpretation of the results was done as described in CLSI 2013[13]. Antibiotic discs (HiMedia Laboratories Pvt. Ltd., India) used were, Cefotaxime (CTX) (30μg), Imipenem (10μg), Piperacillin/Tazobactam (PTZ), and Clavulanic acid (CEC). Control strains of *P. aeruginosa* ATCC 27853 and *E. coli* ATCC 25922 were used in parallel as a part of quality control. Organisms resistant to two or more classes of antimicrobial agents were considered to be multidrug resistance (MDR). The procedure involves nutrient broth having isolated organism and after some time sterile cotton swab was dipped into the suspension. The cotton swab was revolved few times and pressed against the inside wall of the tube and inoculated on the fresh surface of a Mueller-Hinton agar (MHA) plate by streaking the swab over it. For obtaining even distribution, the swab was streaked two or three times at an angle over the MH agar. After 2 minutes, antibiotic discs were applied and pressed down to ensure attachment with agar surface. The plates are then inverted and incubated aerobically at 37°C after the application of disc

### 2.4 Screening of ESBL-Producing Strains

According to CLSI guidelines, strains showing zone of inhibition of ≤25mm for cefotaxime were considered for conformational test for ESBL. ESBL production among potential ESBL-producing isolates was confirmed phenotypically using combined disc method. Comparison of the zone of inhibition was made for the cefotaxime (30µg) discs alone vs. that of the ceftazidime/Clavulanic acid (CAC) (10μg) when placed 25mm apart (center to center). Isolates showing an increase in zone diameter of ≥5 mm around either of the clavulanate combined discs compared to that of the disc alone was considered ESBL producer.

### 2.5 Molecular detection of β-lactamase genes

All the phenotypic ESBL Escherichia coli isolates were subjected to molecular analysis for the confirmation of ESBL production. Molecular detection of Escherichia coli harboring ESBL genes (bla-CTX-M, bla-TEM, and bla-SHV) was carried out by conventional polymerase chain reaction (PCR) with PCR master mix (DreamTaq Green PCR Master Mix; Thermo Scientific) using specific primers (Table 1).

**Table 1:** Total Number of Isolates from Clinical Specimens

| Identified Microorganisms | Number of Isolates | Percentage (%) |
| --- | --- | --- |
| *E.coli* | 50 | 71.4 |
| *Klebsiella sp.* | 14 | 20 |
| *Pseudomonas sp.* | 3 | 4.2 |
| *Citrobacter sp.* | 2 | 2.8 |
| *Shigella* | 1 | 1.4 |

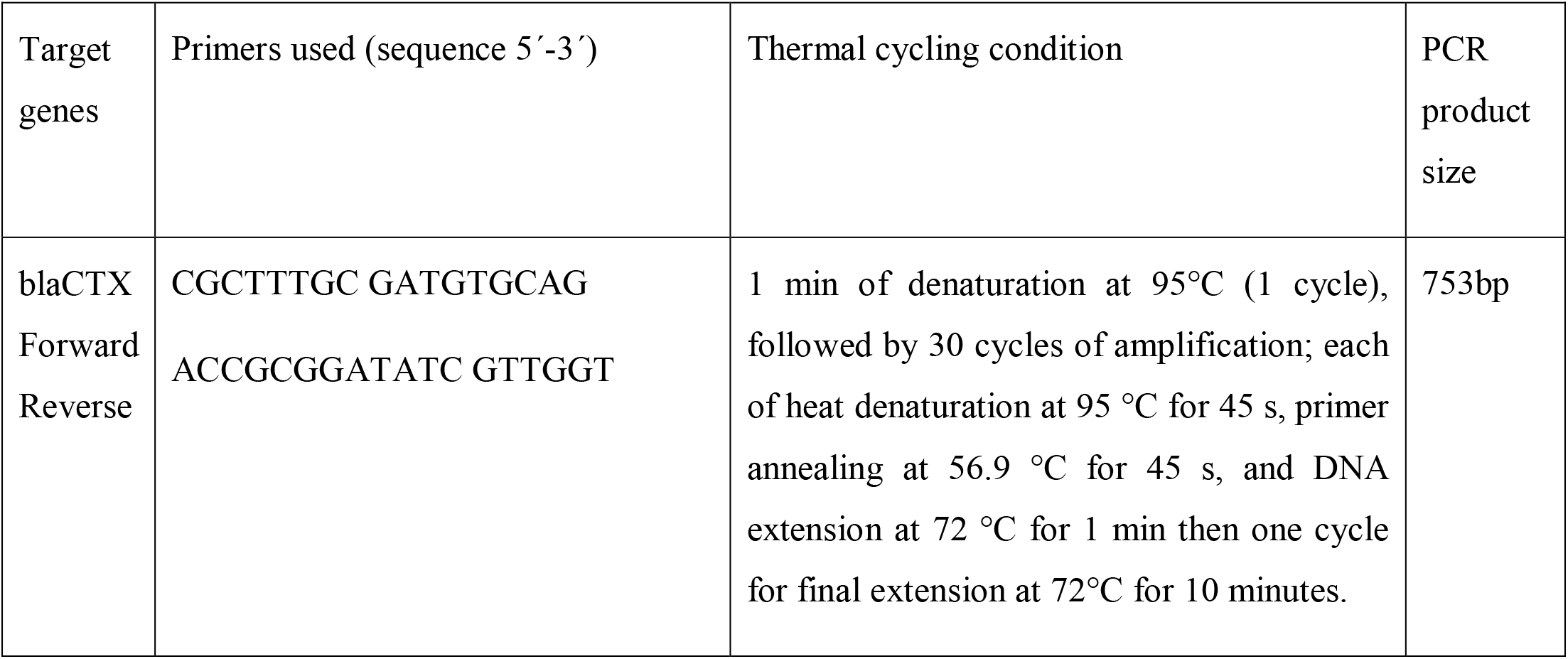
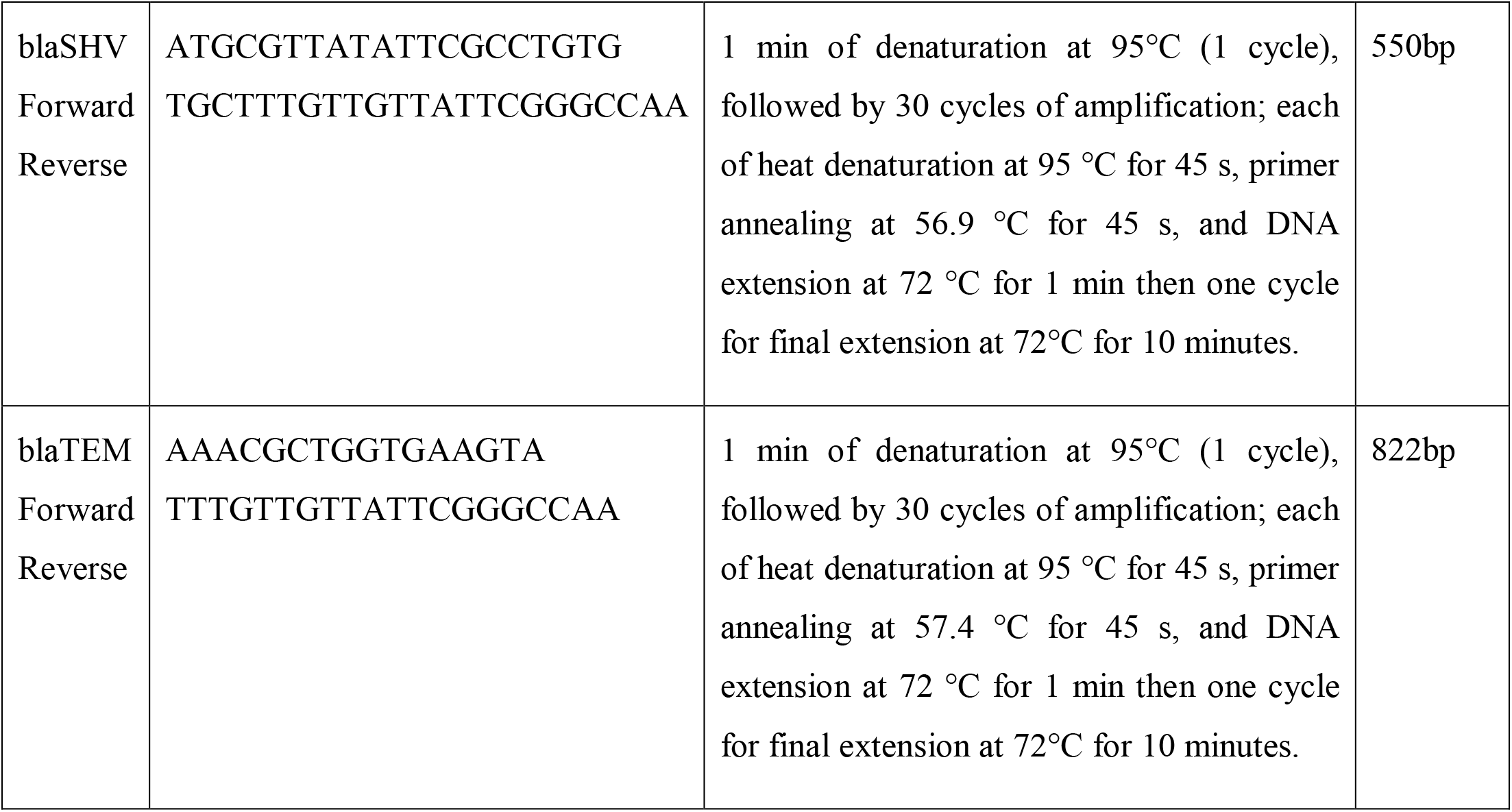

#### Plasmid DNA Extraction and Amplification

For plasmid DNA extraction, a single colony of each potential ESBL-producing isolates was inoculated into Luria-Bertani broth and incubated till the logarithmic state. Extraction and purification of plasmid DNA of bacteria was carried out using a commercial kit (SureSpin Plasmid Mini kit from Genetix Biotech, Asia) following manufacturer’s instructions. Purified DNA from bacterial isolates was used as a template to detect ESBL genotypes: CTX-M, TEM, and SHV β-lactamase genes. Primers for the amplification of ESBL genotypes (bla-CTX-M, bla-TEM, and bla-SHV) were purchased and are as listed in above Table 1.

Polymerase chain reaction-(PCR-) based amplification of ESBL genes was carried out as the method previously described [14]. Amplification reactions were carried out in a DNA thermal cycler (CG) with the following thermal and cycling conditions: initial denaturation at 94°C for 5 min, followed by 35 cycles at 94°C for 30 sec, 45°C for 1 min, and 72°C for 1 min, and a final extension at 72°C for 10 min. The amplified PCR products were subjected to electrophoresis on a 1.5% agarose gel in 0.5 × TBE buffer. DNA was stained with ethidium bromide (1 μg/ml), and the amplified DNA bands were visualized using a UV-transilluminator.

### 2.6 Statistical Analysis

Data were analyzed using IBM-SPSS version 22 (Chicago, USA, 2013). Qualitative data were expressed as number and percentage and quantitative data were expressed as mean ± standard deviation (SD). Chi square test was used to compare frequencies of qualitative data and Student’s t test was used to compare means in quantitative data. P values < 0.05 were considered significant.

## 3. RESULTS

A total of 100 urine and vaginal samples were collected from patients that had symptoms suggestive of UTI. The age of patients ranged from 2 - 80 years (mean 41 years). These patients were 38 males and 62 females. The percentage of women was higher than that of men.

Most of the patients (39%) reporting UTIs were between 21-40 years of age. 31 % patients were 41-80 years old, 26 % patients were above 60 years old and the lowest was 4% patient of age group less than 20.

### 3.1 Distribution of Isolates from Clinical Specimens

One hundred clinical samples were collected from patients (62 females and 38 males) during the study period. The samples obtained were urine (*n* = 77) and high vaginal swabs (*n* = 23). Total isolates obtained was 70. *Escherichia coli* was isolated in the highest number from urine samples. *E.coli* (n=50; 71.4%) and followed by *K. pneumoniae* (*n* = 14; 20%) was a predominate isolate and the rest 8% included *pseudomonas sp*., *Citrobacter sp.and Shigella sp*. (Table 1 and Figure 3).

**Figure 1:**
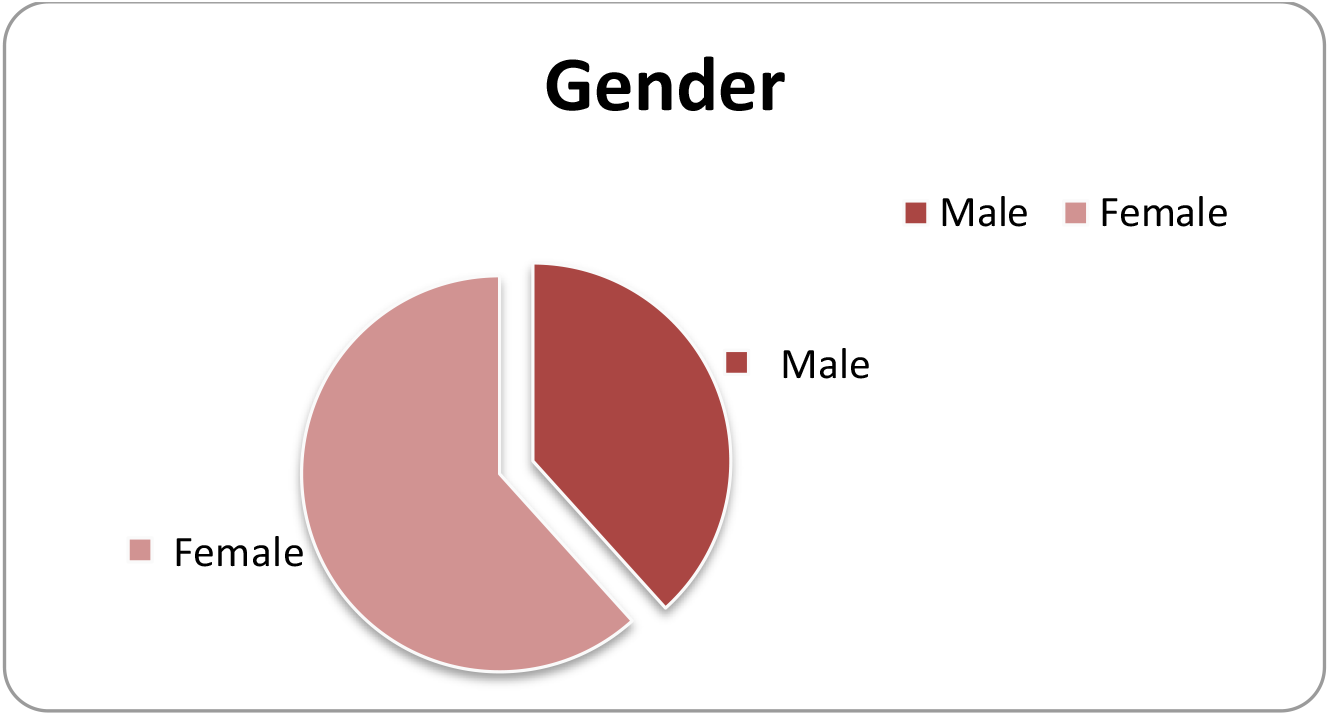
Gender distribution.

**Figure 2:**
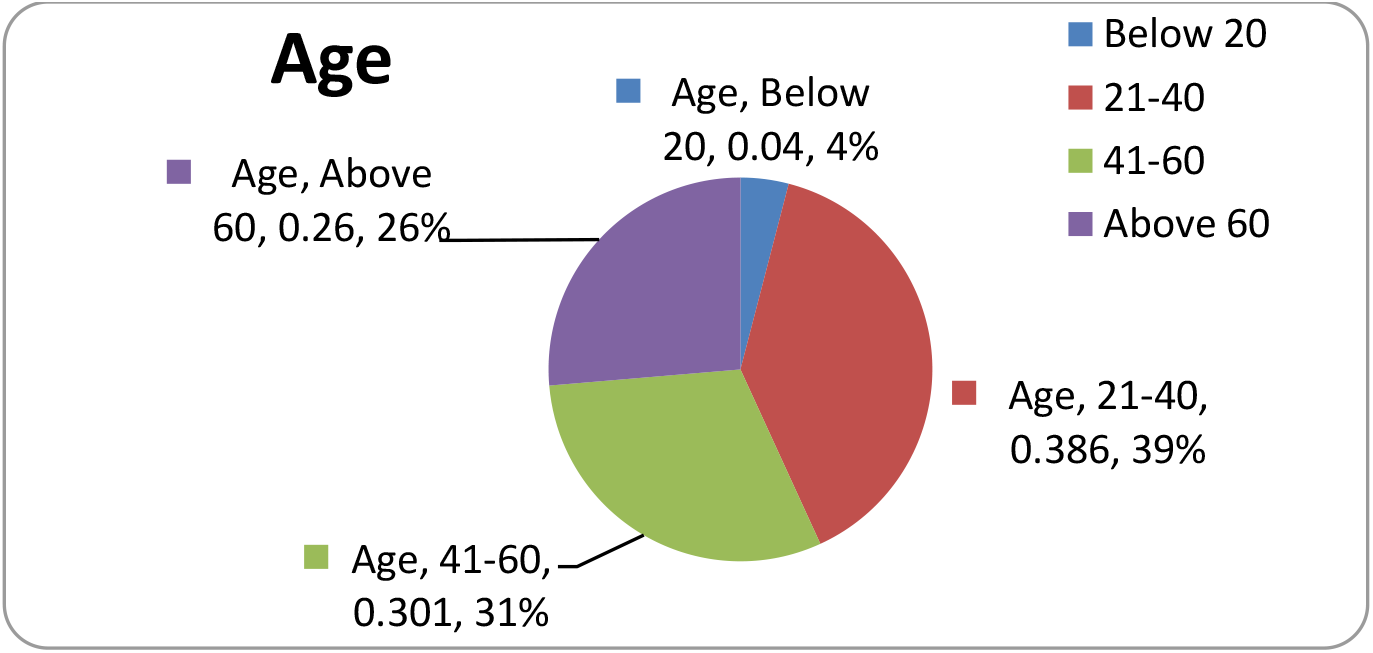
Age distribution.

**Figure 3:**
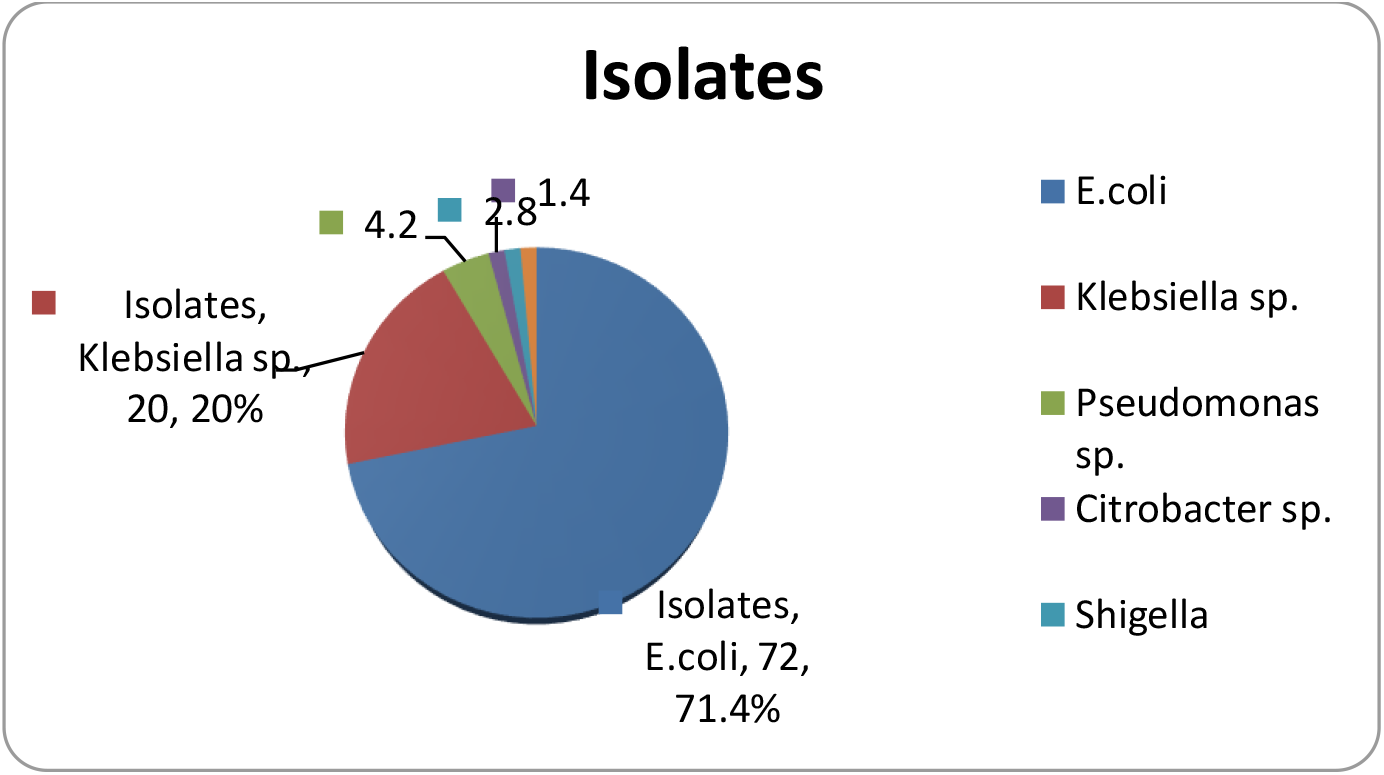
Percentage of patients with *E. coli* infection and non-*E.coli* infection among the isolates.

**Figure 4:**
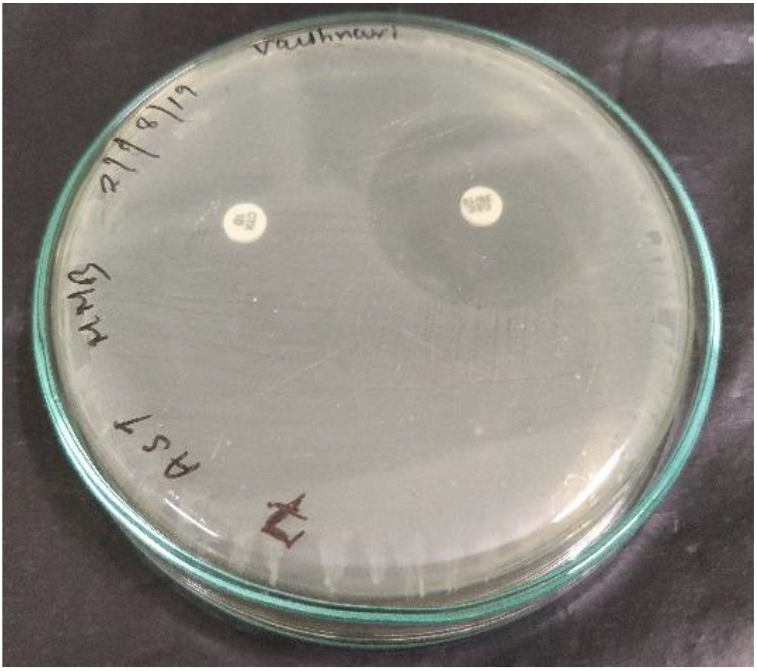
ESBL positive strain.

**Figure 5:**
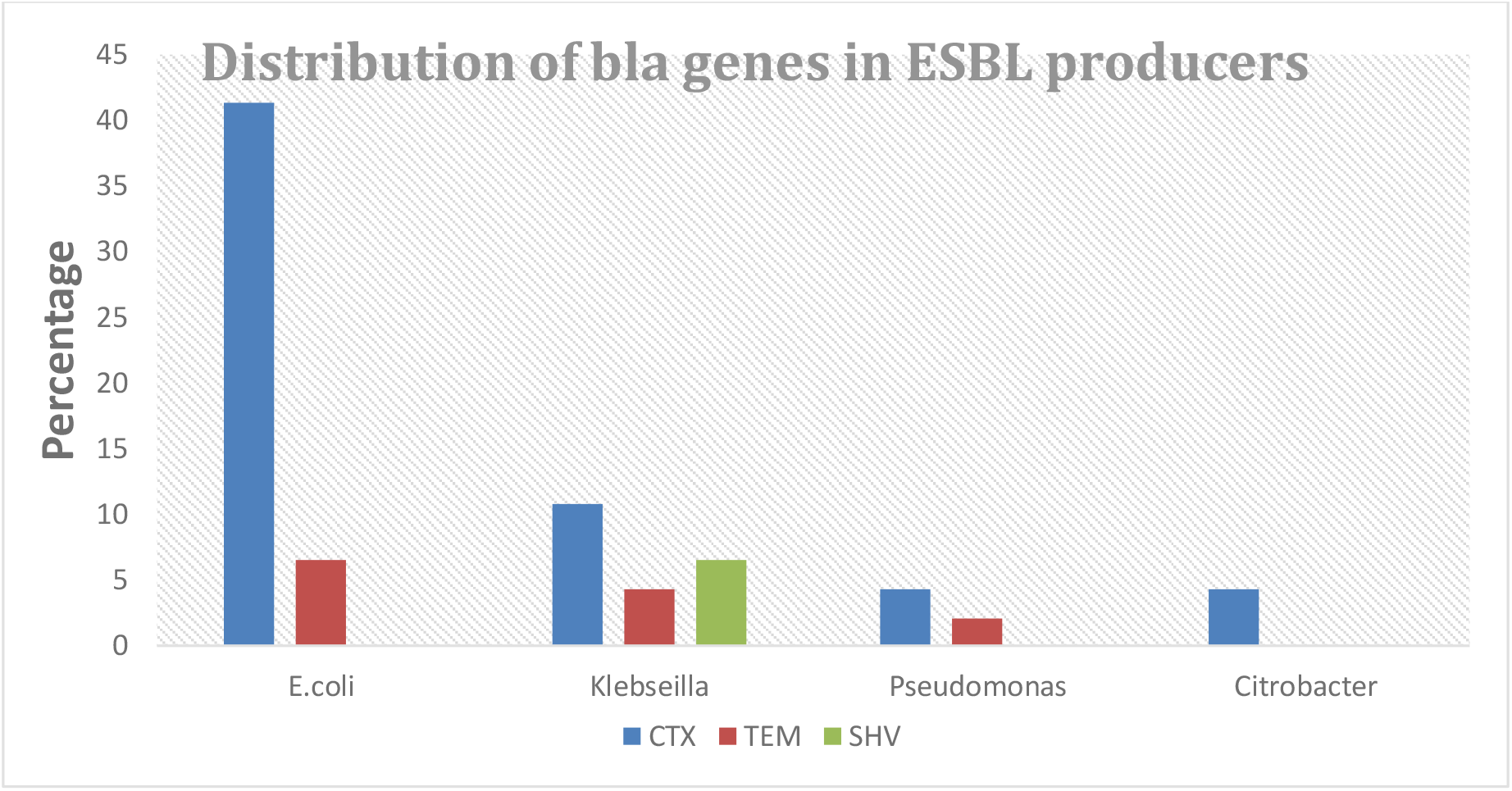
Distribution of bla gene variants in ESBL producing organisms.

### 3.2 Antimicrobial Sensitivity Pattern of Uropathogenic E. coli Isolates

The antibacterial susceptibility profile was examined in all 100 uropathogenic isolates. The resistance rate to each antibiotic was calculated as the number of resistant isolates divided by the total number of isolates. All ESBL-producing isolates showed 100% resistance to ampicillin, and to all cephalosporins including **Cefotaxime** and **Clavulanic acid**. Among the β-lactam/β-lactamase inhibitor combinations 99% were resistant to amoxicillin/clavulanic acid. Out of 70 isolates that were considered as potential ESBL producers, 46 (65.7%) were confirmed as ESBL producers.

### 3.3 Molecular analysis of ESBL genes

Among the 46 ESBL producing bacterial isolates representing *E.Coli, Klebsiella, Pseudomonas, Citrobacter*, were selected and characterized with PCR for genes encoding resistance. Of these, the higher resistance (n=28, 60.8 %) was encoded by *bla*_*CTX*_ genes. *bla*_*TEM*_ and *bla*_*SHV*_ genes were detected in 9 and 19.5 % of the isolates, respectively. 28 were positive for the *bla*_CTX-M_ gene (60.8%), of which 19 were *E. coli*, 5 were *Klebsiella sp*., and 2 each of *Citrobacter sp. and Pseudomonas sp*. There was a small proportion of other gene observed in ESBL producers such as *bla*_*TEM*_ and *bla*_*SHV*_ genes. Out of total 46 ESBL producers, 6 were positive for the *bla*_*TEM*_ gene (13 %), of which 3 were *E.coli* and 1 each of *Klebseilla sp*., *Citrobacter sp.and Pseudomonas sp*. Also, 3 out of 46 ESBL producers showed presence of *bla*_*SHV*_ gene (6.5%) in *Klebseilla sp*. Three (*E. coli*) isolate produced two ESBLs (TEM and CTX) and Klebsiella isolate produced all types of ESBLs (CTX, TEM & SHV). 37.3% of bacteria harbored multiple *bla* genes (≥ two genes) (Table 2). All ESBLs were resistant to cefotaxime..

**Table 2:** Distribution of bla gene variants in ESBL producing organisms.

| Isolates (46) | *CTX* | *TEM* | *SHV* |
| --- | --- | --- | --- |
| *E.coli* | 19 | 3 |  |
| *Klebseilla sp.* | 5 | 2 | 3 |
| *Pseudomonas sp.* | 2 | 1 | - |
| *Citrobacter* | 2 | - | - |
| Total | 28 | 6 | 3 |

## 4. DISCUSSION

In this study, we investigated the antibiotic resistance profiles and trends in MDR isolates obtained from urine and vaginal samples. Over the past two decades, there has been a significant number of infections caused by bacteria expressing extended-spectrum-β-lactamase (ESBL) and carbapenemases [15]. It has been frequently reported that UTIs occur far more frequently in girls than in boys during the first few months of life, presumably due to the shorter length of the female urethra[16].

As expected, 62% of isolates out of 100 were that of women which is 3/5^th^ of total isolates. ESBL mediated resistance was found in 65.6% of our isolates out of 70 potential ESBL producers. This prevalence rate is much higher than other reports from India and abroad. However, the prevalence of ESBL producing strains *of E.coli, pseudomonas sp*. and members of *enterobacteria* have to be carefully monitored to prevent misuse and overuse of antibiotics because antibiotic exposure could make them more vulnerable to *multidrug resistant bacteria*.

The prevalence of ESBL production varies according to species, geographical areas, variations in infection control programs, different patterns of empiric antibiotic regimens and even over time. Moreover, selective pressure caused by the overuse of cephalosporins in some countries leads to the emergence of increasing rates of ESBLs production[17].

Previous report from North East India showed high rate of *bla*_SHV_ (63.4%) followed by *bla*_CTXM_ (60.86%) and *bla*_TEM_ (54.3%). A study from Central India (Rajasthan) on 20 ESBL-producing *E. coli* isolates reported a high rate of *bla*_CTXM_ (80%) followed by *bla*_TEM_ (60%) and *bla*_SHV_ (55%) [18]. Although a rapid and global spread of CTX-M-type ESBL has been observed since the 2000s, most of the ESBLs recovered in this study accordingly produced a CTX-M-type -lactamase.

The spread of ESBL genes is related to different mobile genetic elements, such as plasmid, transposons and integrons. In our study, Plasmid DNA isolation method was used for extraction of ESBL genes from ESBL producers by Plasmid DNA Extraction kit. The current limited development of novel drugs and substitutes makes the use of ARGs monitoring even more important to develop comprehensive and integrative measures for antimicrobial resistance.

Resistance of *Enterobacteriaceae* to third generation cephalosporins is a worldwide problem [19], which is mainly caused by ESBLs production. Production of additional β-lactamases (AmpC) also contributes to this problem, moreover, the presence of AmpC genes is often associated with multidrug resistance [20]. Previously, AmpC -β-lactamase has received less attention, but is now identified as an important cause of resistance in *Enterobacteriaceae* species. Global spread of β*-*lactamases-producing strains gives a great importance to the study of these strains in community and hospitals for reassessment of the existing treatment protocols. Although MDR rate among ESBL producers in the current study (28.5%) was lower than that reported in previous studies; (96.3%) [21] and (77.6%), there was statistically significant increase in MDR rate reported in the ESBL-producers (28.5%) than that reported in the non-ESBL-producers (1.5%) (*p* value = 0.04).

Out of 72 ESBL-producing isolates in the current study, 48 (99%) isolates were positive for ESBL genes, with *blaCTX-M* type as the most predominant. The frequency of community-acquired infections caused by *blaCTX-M*-producing strains have markedly increased in the last decade, that agrees with our findings, where *blaCTX-M* genes were detected in 254 (81.6%) of *Enterobacteriaceae* isolates. Our results concur with several studies on hospital and community-acquired infections, those reported high prevalence of *blaCTX* among *Enterobacteriaceae* species in Egypt, Burkina Faso, Iran, Qatar and Japan. *bla*_*TEM*_ and *bla*_*SHV*_-producing strains were reported previously as hospital pathogens until the late 1990s [22].

## 6. CONCLUSION

High burden of antimicrobial resistance and increased prevalence of ESBL-producing Escherichia coli associated with UTI are the major findings of this study. Diverse genotypes of ESBL E. coli along with resistance towards common antibiotics were observed. The *bla*_*CTX-M*_ type was the pre-dominant among ESBL-producing Enterobacteriaceae, especially in combination with *bla*_*TEM*_ enzymes. Molecular detection and identification of beta lactamases is essential for a reliable epidemiological investigation of resistance. These enzymes can be chromosomal or plasmid mediated, which may help in the distribution of antimicrobial drug resistance in health care organization. In this perspective, regular national-wide epidemiological surveillance of bacterial pathogens causing UTIs and their anti-microbial resistance would be useful in developing the treatment guidelines in our country. Further next generation sequencing (NGS) studies are essential for a more comprehensive analysis of ESBL form.

## Financial support & Sponsorhip

We are very grateful to IRSC, Dr. B. Lal Institute of Biotechnology for their kind funding support.

## Conflicts of interest

**NIL**

## Notes

### Competing Interest Statement

The authors have declared no competing interest.

